# Image-centric compression of protein structures improves space savings

**DOI:** 10.1101/2022.01.20.477098

**Authors:** Luke Staniscia, Yun William Yu

## Abstract

**Background:** Because of the rapid generation of data, the study of compression algorithms to reduce storage and transmission costs is important to bioinformaticians. Much of the focus has been on sequence data, including both genomes and protein amino acid sequences stored in FASTA files. Current standard practice is to use an ordinary lossless compressor such as gzip on a sequential list of atomic coordinates, but this approach expends bits on saving an arbitrary ordering of atoms, and it also prevents reordering the atoms for compressibility. The standard MMTF and BCIF file formats extend this approach with custom encoding of the coordinates. However, the brand new Foldcomp tool introduces a new paradigm of compressing local angles, to great effect. In this article, we explore a different paradigm, showing for the first time that image-based compression using global angles can also significantly improve compression ratios. To this end, we implement a prototype compressor ‘PIC’, specialized for point clouds of atom coordinates contained in PDB and mmCIF files. PIC maps the 3D data to a 2D 8-bit greyscale image and leverages the well developed PNG image compressor to minimize the size of the resulting image, forming the compressed file.

**Results:** PIC outperforms gzip in terms of compression ratio on proteins over 20,000 atoms in size, with a savings over gzip of up to 37.4% on the proteins compressed. In addition, PIC’s compression ratio increases with protein size.

**Conclusion:** Image-centric compression as demonstrated by our prototype PIC provides a potential means of constructing 3D structure-aware protein compression software, though future work would be necessary to make this practical.

## 1 Background

For over half a century, determining protein structure has been a primary means of understanding function and behavior [1, 2]. After proteins are characterized by researchers using various methods such as X-ray crystallography, NMR spectroscopy, and cryo-electron microscopy, various files are generated describing the protein and stored in online repositories such as the Protein Data Bank [3, 4]. One such file, the FASTA file, contains strings of characters representing the amino acids that make up the protein and its variants [5]. Other files, such as PDB (Protein Data Bank format) and mmCIF (macromolecular Crystallographic Information File) files, contain structural information about the protein [6]. Although the Protein Data Bank is no longer growing exponentially, the number of new structures deposited is still quite formidable [4]; furthermore, the recent publication of AlphaFold predicted structures has increased total available structures by orders of magnitude [7].

FASTA files are used for storing both protein and genomic sequence information, and much work has been done to create customized sequence compression algorithms. It bears mentioning that the genomic sequence compression literature has recently seen significantly more activity with the advent of next-generation sequencing [8, 9, 10, 11], and many protein sequence compressors take advantage of that work. For protein sequences, [12] introduce a single and double pass version of a amino acid sequence compressor for FASTA files that makes use of substitution matrices. MFCompress was introduced by [13], and converts the amino acid sequences to their corresponding DNA bases, divides the data into three streams, and compresses the resulting streams. CoMSA is another compression algorithm for FASTA files introduced by [14] based on a generalized Burrows-Wheeler transform. Similarly to MFCompress, The Nucleotide Archival Format (NAF) introduced by [15] is another compressor that works on amino acid sequences converted to their corresponding DNA bases by dictionary encoding this transformed string.

In addition to directly transforming and compressing the sequences in FASTA files, a significant amount of research has gone into read-reordering algorithms for genomic sequences in the BEETL [16], SCALCE, [17], MINCE [18], and more. These methods are applicable when FASTA (and the related FASTQ) files are used to store multiple small fragments (‘reads’) of sequences; next-generation sequencing produces these reads in no particular order, so the reads can be safely reordered without losing important information. When properly performed, this reordering can significantly improve the compression ratio of standard compressors.

On the other hand, the primary data component of PDB and mmCIF protein structure files is a point cloud of coordinates belonging to the atoms that make up the protein. In the standard formats, each atom has its own separate ASCII-formatted line entry in the file that contains the type of atom, type of amino acid to which it belongs, atom and amino acid identifiers, followed by three floating point Cartesian coordinates, along with other information. The coordinates are measured in units of Angstroms *Å*, where 1*μm* = 10, 000*Å* [19]. Unlike their FASTA counterparts, comparatively less work has been done to create compressors customized for the structural data contained in PDB and mmCIF files, though there have been a number of recent tools/formats like MMTF [20], BCIF [21], and the brand new Foldcomp [22]. We note with especial interest Foldcomp, which introduces a new paradigm for compressing atomic coordinates using local angles, which is a radical shift from what both MMTF and BCIF do. In this manuscript, we explore yet another different direction in the form of image-centric compression and global angle computations.

[23] did a deep investigation on compressing 3D coordinates of atoms in proteins by investigating a full gamut of compression techniques. Their final recommendation was to apply “intramolecular compression”, which aims to reduce the size of each protein via three steps: encoding, packing, and entropy compression. The encoding step transforms floating point coordinates into alternate representations, such as Integer, Delta, Predictive, Wavelet, and Unit Vector encodings. Integer encoding as described by [23] multiplies the floating point location coordinates by a power of 10 and rounds the result to the nearest integer. This encoding strategy is lossy when not all decimal places of precision are kept in the integer encoded value, but it can be lossless when used in MMTF and BCIF with a sufficiently large power of 10. However, some amount of loss of precision can be acceptable because of both measurement error, and due to the natural uncertainty of exact atom locations in a protein—PIC will use a lossy variant of integer encoding. Going back to [23], the authors suggest that after packing the encoded coordinate vectors using either recursive indexing or variable packing, the resulting packed coordinates are entropy encoded using standard methods like gzip [24] or brotli [25], which are both combinations of LZ77 dictionary based encoding and Huffman encoding.

However, [23]’s investigation focused primarily on compression of atomic coordinates as sequential objects stored within a text file, treating the data as sequential, much like in FASTA files without reordering. However, unlike protein/genomic sequences, 3D atomic point clouds are not naturally sequential, and the sequence of atoms listed is purely an artifact introduced by using a sequential file format to store the atoms. Thus, preserving the order of the atoms as listed out in a PDB or mmCIF file is largely irrelevant for the purposes of compression, so long as the original information can be reconstructed. Given the background above, one logical next step would be to perform a principled reordering of the atoms to improve compressibility, similar to the technique used by read-reordering algorithms (where again, the order of reads output by the sequencer is inconsequential). The remaining question is of course how to perform that reordering, as point clouds are very different from sequenced genomic reads in underlying structure.

To resolve this question, we turn to an alternate paradigm for compressing point cloud data sets proposed in the field of LIDAR (light detection and ranging) imaging. [26] proposed a new compression algorithm for the 5D point cloud data generated by LIDAR scans of real-world scenes. The LIDAR scans produced tuples of data points containing coordinates of a point in space in the scene, along with reflectance and colour data of the surface at that location. Their compression algorithm converts the Cartesian coordinates to spherical coordinates, maps the angular coordinates to the axes of an image, and the radial component, colour, and reflectance data to pixel’s fields at the mapped location. The radial component, colour, and reflectance data are written to the R, G, and B components respectively of a single coloured image as well as the greyscale intensity field of three separate consecutive images. The resulting images were compressed using PNG, JPEG 100 (lossless, perfect quality JPEG), JPEG2000, no compression TIFF, LZW TIFF, and Pack Bits TIFF lossless image compressors. The authors of [26] found that compressing three greyscale images using the PNG compressor performed the best in terms of compression ratio.

In this manuscript, we take inspiration from the next-generation sequencing read-reordering literature and combine the intramolecular compression techniques of [23] with the image-centric methods of [26]. In section 2, we outline our new compression algorithm, PIC, for the structural protein data contained in PDB and mmCIF files. Design choices and methodology are examined in detail followed by a pseudo-code outline of the compression algorithm. In section 3, we give compression results for the atomic coordinates of 20 proteins of a variety of different sizes compressed using both PIC and gzip and show PIC outperforms gzip in terms of compression ratio for proteins over 20,000 atoms in size. We also give the images that constitute the compressed files for a few of the compressed proteins. Furthermore, although PIC is not a full compressor as it does not compress metadata, for the sake of completeness, we also compare PIC file sizes against full compression software MMTF, BCIF, and Foldcomp. In section 4, we highlight some trends in the compression results and make note of the advantages of the PIC compressor over gzip for structural protein data compression.

### Algorithm 1 PIC compression algorithm

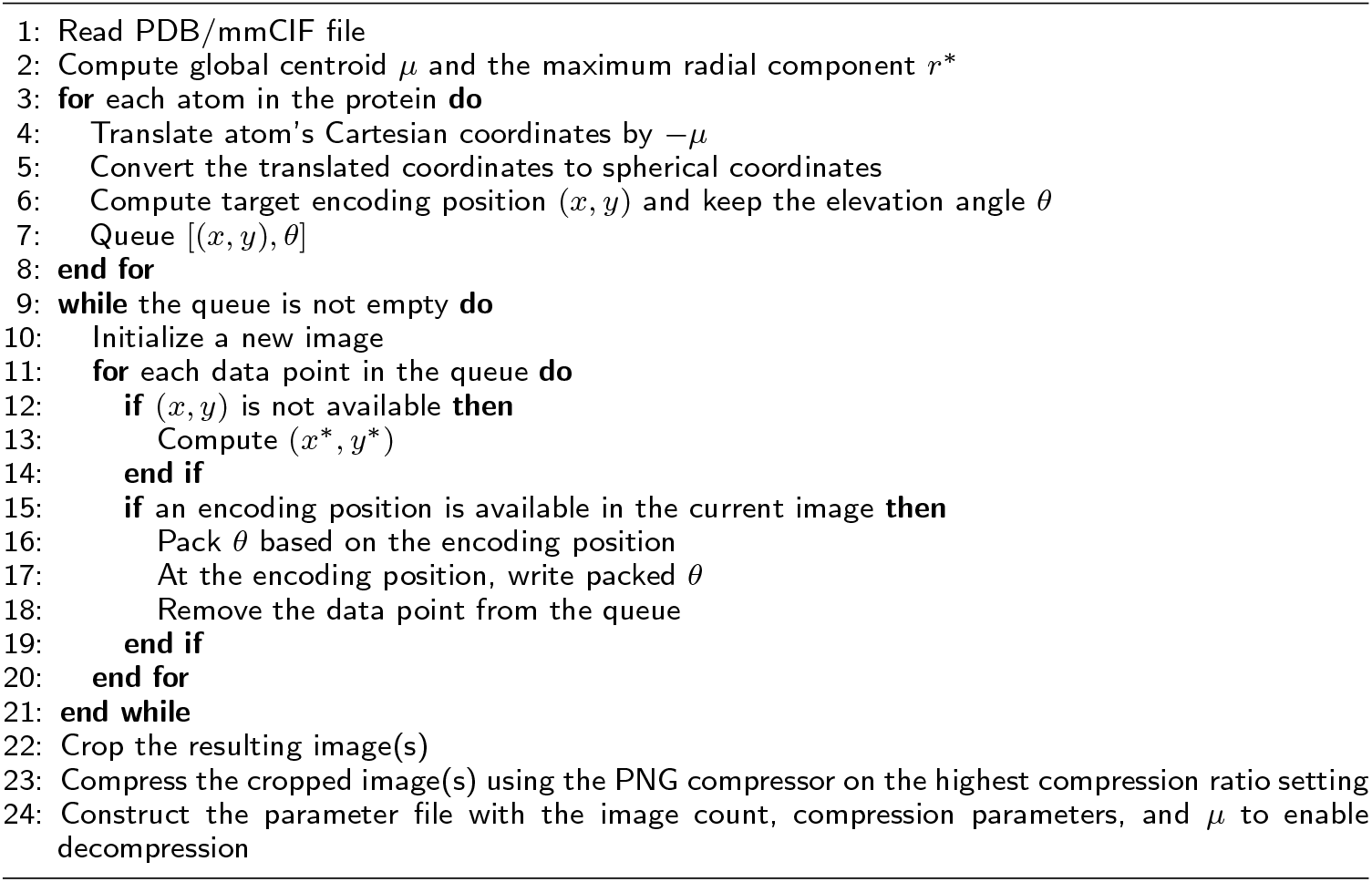

## 2 Implementation

The PIC compression algorithm has has three main components, namely mapping each atom to a position in an image, encoding information at that position, and compressing the resulting image. A high-level overview is given in algorithm 1 and figure 1, and details are furnished in the following text.

**Figure 1.**
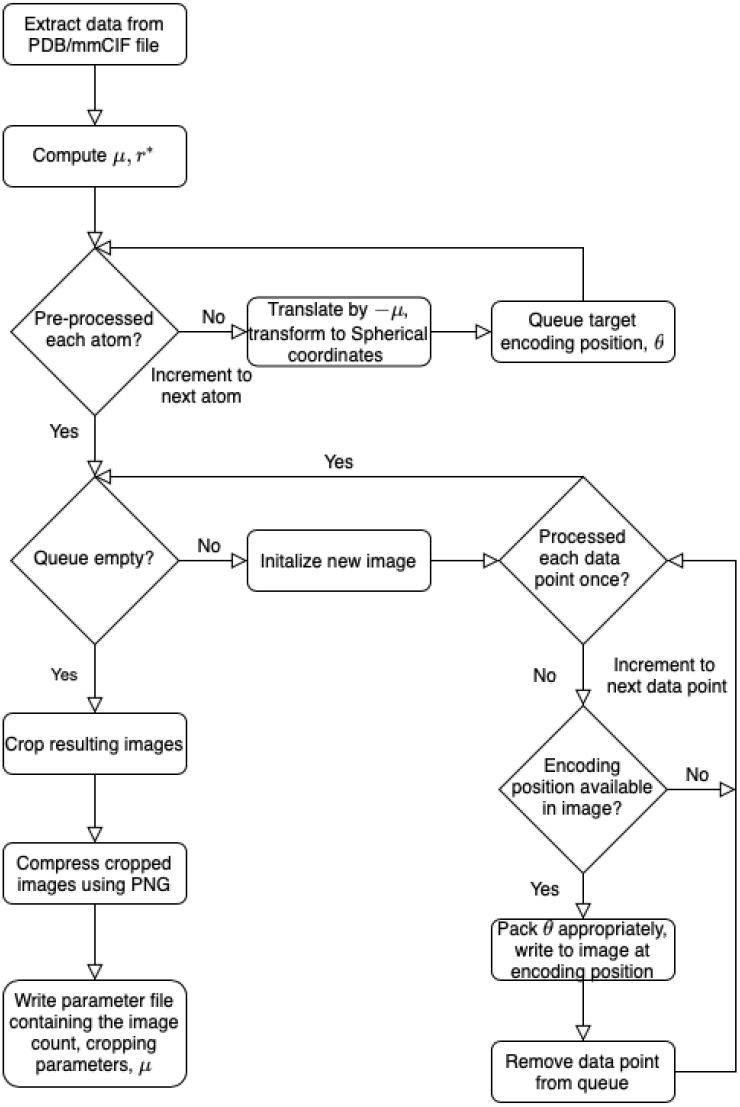
Flow chart diagram of the PIC compression algorithm. *μ* is the global centroid of all atoms used to center the image, and *r*\* is the maximal radial component that needs to be stored after centering. The basic intuition is to store atoms and their coordinate data in a pixel corresponding to the radial coordinates, and then compress with PNG.

### 2.1 Mapping

Cartesian coordinates of atoms stored in the protein’s PDB or mmCIF file are extracted and the global centroid *μ* of all the coordinates is computed. The coordinates are translated by *−μ* so that the global centroid becomes the new reference point or origin for the coordinates. This transformation minimizes the instances of collisions when mapping the coordinates to the image. To decompress the images, *μ* is stored along with the images.

The translated coordinates are then transformed to spherical coordinates. Each spherical coordinate component is rounded to a precision of one decimal place. [23] noted that experimental measurements that produce the Cartesian coordinates determine an atom’s position with a degree of uncertainty, greater than 0.2*Å*. This allows for the exploitation of lossy compression to store the coordinates only up to a tenth of an *Å*, which is generally sufficient to preserve the essential structural information provided by lossless representation.

The radial *r* and azimuth *φ* spherical coordinate components of each atom are positionally encoded to the horizontal and vertical axis of an eight bit pixel greyscale image as follows

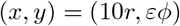

where *ε* is a user-defined parameter that sets the number of pixels per azimuth angle degree. Letting *r*\* be the maximal radial component across all spherical coordinates, the dimensions of the resulting image are 10*r*\* *×* 360*ε*. Further note that while *x* ∈ ℤ_*≥*0_, *y* is not necessarily an integer. However, *ε* is chosen such that 360*ε*, 8*y* ∈ ℤ_*≥*0_ for all *y* and *ε ≥* 1.25. This ensures there is at least one bit available per tenth of an azimuth angle degree and each *y* coordinate has an integer bit-level position on the vertical axis. In this way, we view each column in the image as a bit sting that is being written to.

Care must be taken when choosing *ε*. Setting *ε* too large will produce a large image, degrading the compression ratio. On the contrary, choosing a small *ε* will induce more collisions when data is mapped to the image. This results in increased compression time, as alternate data storage locations need to be considered. A decrease in the compression ratio may also be experienced in this case as more data points will need pointers to their intended locations and additional images may need to be populated to store all the required data.

The remaining elevation angle *θ* is stored in the image’s pixel intensity values beginning at the data point’s (*x, y*) encoding position in the image. Further details on how the elevation angle is formatted or *packed* and stored in the image is described in section 2.2. This encoding scheme was selected as it positionally encodes the spherical coordinates *r* and *φ* with the largest range of values and encodes the smallest ranging coordinate *θ* in the image’s pixel’s intensity values. Thus each coordinate takes up the fewest amount of pixels when encoded into the image, allowing for more data to be stored in the image before another image needs to be generated.

In the event that a data point is mapped to a position that does not have availability to hold all the required information, an alternate encoding position (*x*\*, *y*\*) is determined systematically. A position (*x, y*) has availability if all bits at positions between and including (*x, y*) and (*x, y* + *l −* 1*/*8), where *l* is the length of the encoded elevation angle in bytes, have not had data previously written to them. Beginning at the data point’s target encoding position (*x, y*), the positions (*x, y* + *i/*8 mod 360*ε*), 0 *< i <* 8 *·* 360*ε* = 2880*ε* are scanned subsequently to find the first position with availability. This position is the alternate encoding position. All encoding positions (*x, y*) also satisfy *y < y* + *l −* 1*/*8 *<* 360*ε*, ensuring no data points begin at the bottom of the image and finish at the top to enable proper decompression of the image.

If (*x*\*, *y*\*) is the alternate encoding position for a data point with target position (*x, y*), and *y ≤ y*\* *< y* + 0.1*ε*, the encoded elevation angle is stored begining at (*x*\*, *y*\*) as is. Otherwise, a pointer *p* is encoded and stored along with the encoded elevation angle at (*x*\*, *y*\*). *p* points to the largest *y*^′^ ∈ {*i/8|0 ≤ i <* 2880*ε*} that satisfies *y ≤ y* ^*′*^ *< y* + 0.1*ε*, namely *y*^*′*^ = *y* + 0.1*ε −* 1*/*8. The stored pointer is the integer *p* = 8(*y*\* *− y*^*′*^). Note that *p >* 0 as *y*\* *> y* ^*′*.^ The decompressor then knows that the intended azimuth angle for the data point is that belonging to the position (*y*\* *− p/*8) = *y* ′.

In the case that an alternate encoding position cannot be found in the current image, another image is generated, if not already done by a previous data point. The above mapping procedure is repeated in that image to locate an encoding position for the data point. This process repeats until an encoding position is determined for each atom’s coordinate.

### 2.2 Packing

Elevation angles are stored in pixels’ greyscale values beginning at their corresponding data point’s (*x, y*) encoding position. Each pixel has an 8-bit intensity field. Due to the variable lengths of the binary elevation angles and use of pointers, the following packing scheme is used to store the elevation angles so they can be properly decompressed.

If no pointer is needed, the elevation angle is integer encoded as 10*θ* and converted into its binary representation. If the binary representation has length less than [log_2_(1801)] = 11 bits, 0 bits are added to the front until the representation is 11 bits long. Two additional bits 1 and 0 are added to the front of the resulting binary string in that order to signify the start of a new data point and to notify the decompressor the data point has no pointer, respectively.

If a pointer is required, a similar but expanded packing scheme is used. The second bit is set to 1 instead of 0 to signify to the decompressor that the data point has a pointer. The pointer *p* is converted to its binary representation and prefixed with 0 bits until it has length [log_2_(2880*ε*)]. The adjusted binary representations of the pointer and elevation angles follows the two bit prefix in that order.

For 0 *≤ i <* 8*l*, bit i of the packed string is mapped to the bit at position (*x, y* + *i/*8) in the image. This packing scheme ensures that each data point has one of two possible lengths, the exact length of which can be determined by the second bit located at (*x, y* + 1*/*8). This is a key feature that allows for the proper decompression of the image.

### 2.3 Cropping and Compression

The resulting image(s) are cropped and compressed using the PNG lossless image compressor on the highest compression ratio setting. These image(s) make up the compressed version of the protein’s point cloud of atom coordinates in the PDB or mmCIF file.

Images are cropped to remove any all-black rows and columns on the edge of the image. To decompress the images, two cropping parameters are stored along with each image generated to reverse the cropping.

Other lossless image compressors investigated in [26] were also examined. Similarly to the results found by Houshiar at al., PNG was selected for use in the algorithm as it offers the highest compression ratios of the aforementioned compressors at comparable compression times.

### 2.4 Decompressed File

The original and decompressed files are identical up to the coordinates of the atoms. As noted in section 2.1, since there is a tolerance of up to 0.2*Å* in each coordinate component, each decompressed coordinate is within a euclidean ball of radius 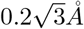 about the original coordinate.

## 3 Results

### 3.1 Atomic cloud coordinate compression

We benchmarked PIC against the gzip compression after the integer compression/precision reduction of [23], the primary relevant prior work. [23] explored a variety of methods for entropy compression, but we found the differences between methods to be swamped out by the integer encoding step, and thus chose gzip as a representative method for sequential compression. We did not compare against plain gzip for that reason, as the compression ratios without the integer compression were not at all comparable. All further references to gzip are to gzip after the [23] integer compression.

Table 1 gives statistics and compression results on 20 proteins compressed using gzip and PIC where *ε* = 2.5 and the decompressed files are identical to the original with the lossy coordinate transform. Figures 2 and 3 compare the 3D structures of proteins to the images created by the PIC compressor. Figure 4 compares PIC vs gzip compression ratios, whereas Figure 5 visualizes some of the same results found in table 1, but plotted against atom size.

**Table 1:**
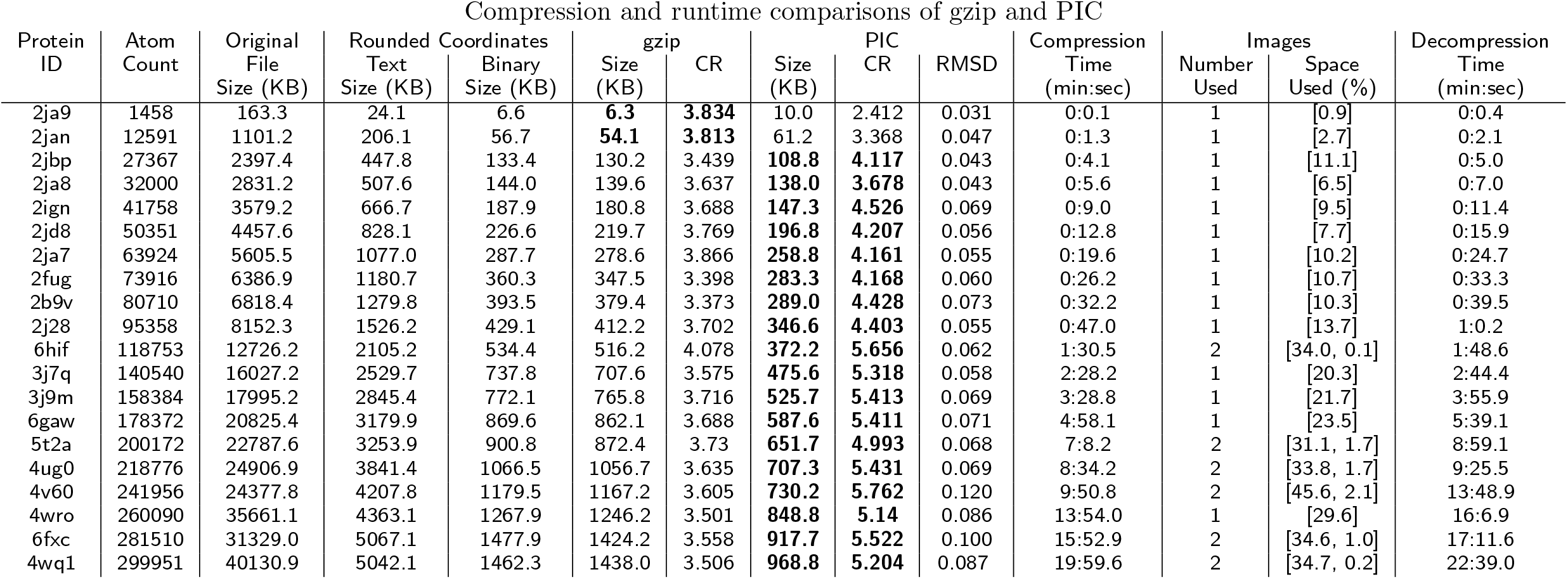
PIC compression algorithm, *ε* = 2.5, results. Rounded Coordinates Text Size and Binary Size are the sizes of the text and binary files (in kilobytes, i.e. 1000 *×* bytes, rather than kibibytes), respectively, that contain only the Cartesian coordinates found in the original file, rounded to one decimal place. The binary file (which uses a variable-length encoding) is then gzipped. The gzip and PIC compression ratios (CR) are the ratios of the Rounded Coordinates Text Size to the size the gzip file and PNG image output(s) from the PIC compressor, respectively. Bolded values are the best of gzip and PIC. Compression and decompression times are for the PIC algorithm; note that our code is unoptimized, as the focus is on compression ratios, but we include these times here for completeness. As an aside, (de)compression for gzip takes negligible time for files of this size. We also include RMSD values to measure the lossiness of PIC compression. Image Space Used gives the proportion of the image space that was used to encode the protein coordinate data, or part thereof, in each image constructed by the PIC compressor (for large proteins, more than one image is needed to represent all the atoms).

**Figure 2.**
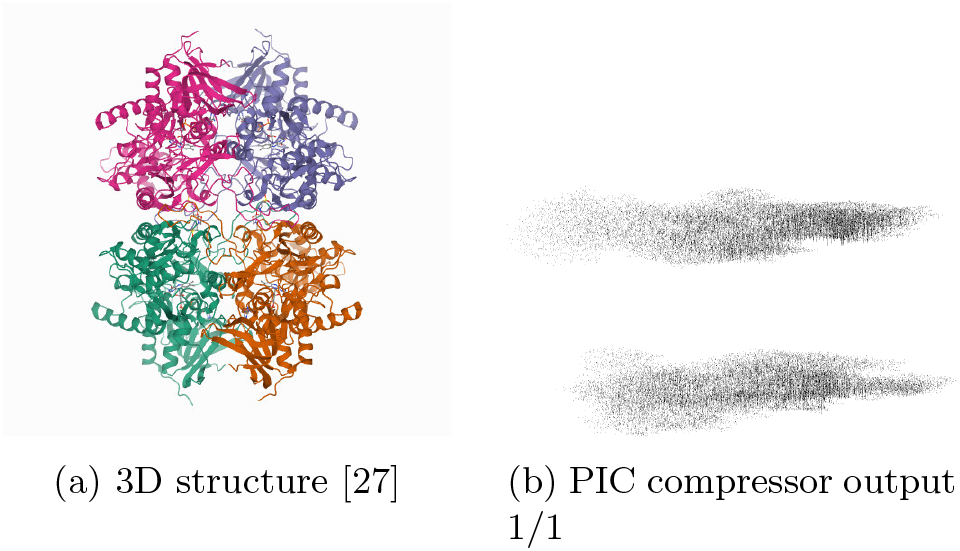
3D structure and PIC compressor PNG image output for 2ign. Some attributes and symmetries in the 3D structure are observed in the corresponding PIC-compressed image. The upper and lower parts of the 3D structure of protein 2ign can be seen in PIC generated image as two separate masses of black pixels, one over the other.

**Figure 3.**
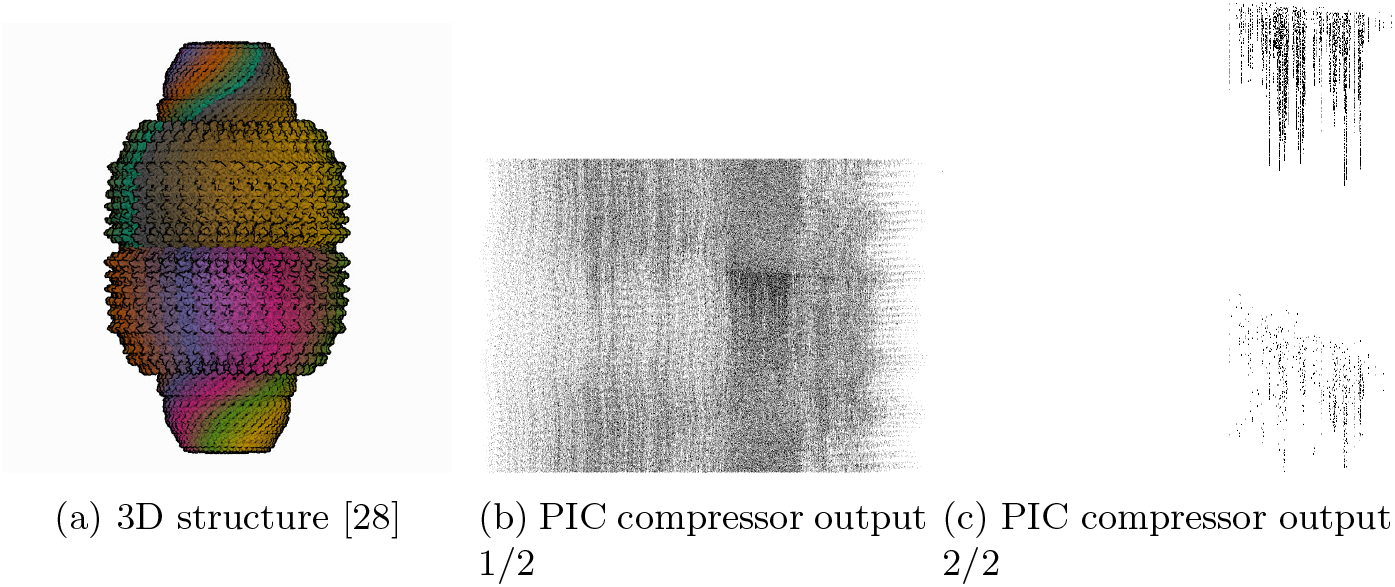
3D structure and PIC compressor PNG image output for 4v60. Some attributes and symmetries in the 3D structure are observed in the corresponding PIC-compressed image. The spiked edge of the 4v60 protein can be seen on the right side of the first outputted image from the PIC compressor.

**Figure 4.**
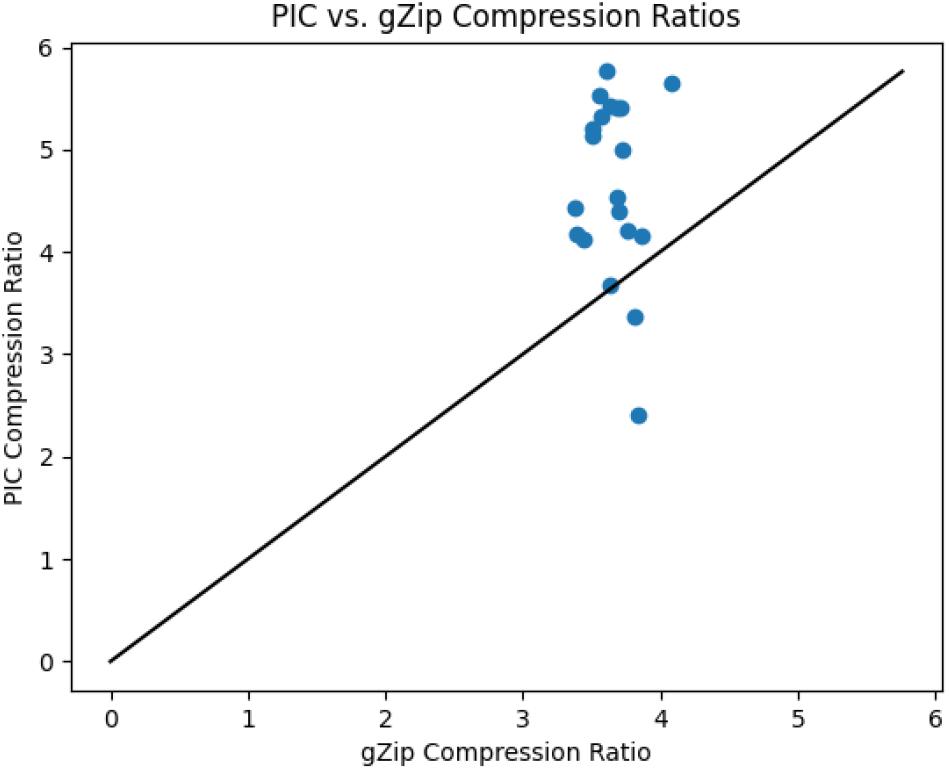
PIC compression ratios plotted against gzip compression ratios (using integer precision reduction for both methods reduced to a tenth of an angstrom for comparability) for each protein compressed in table 1. Points in the region above the diagonal indicates a protein with better compression ratios using PIC than gzip. Vice versa below the diagonal. PIC demonstrates substantially higher compression ratios for nearly all proteins tested.

**Figure 5.**
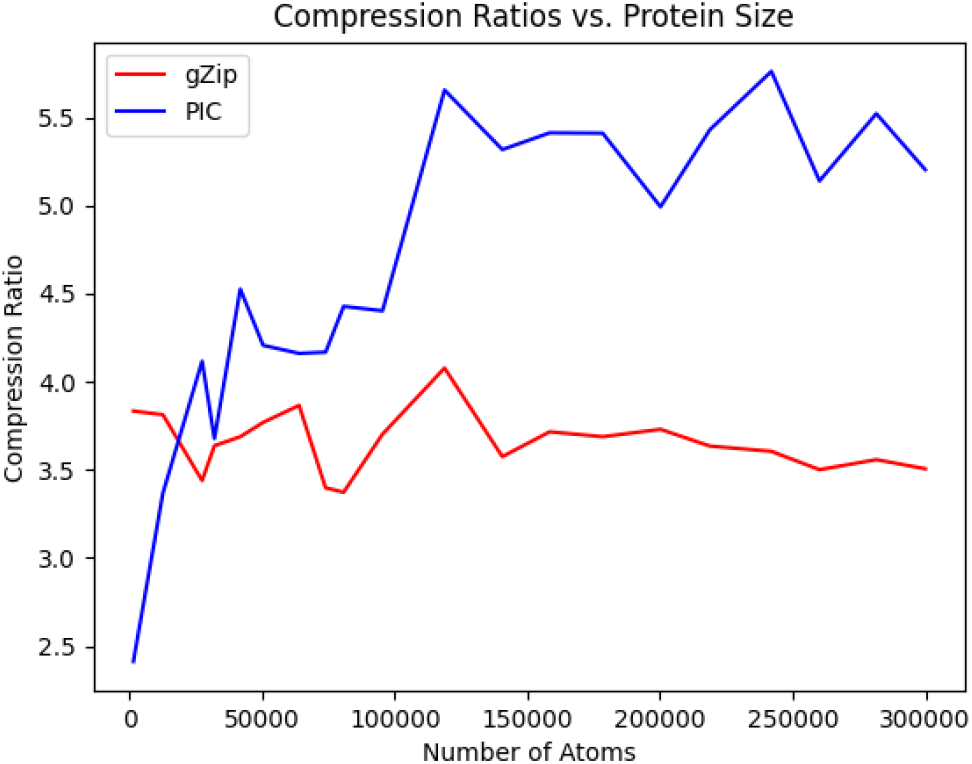
PIC and gzip compression ratios (using integer precision reduction for both methods as suggested by [23] for comparability) for the proteins compressed in table 1 plotted against the number of atoms that make up the compressed protein. All but the smallest proteins showed a higher compression ratio when using PIC; for the small proteins, the extra overhead of PIC dominates, but for any large protein, PIC performs better. Note that this comparison is fair to gzip, as instead of gzipping the original files, we only apply gzip after using the same lossy precision encoding that PIC uses; thus, the comparison here is really between sequential storage of a text file using gzip, and spherical storage using PIC.

As can be seen from table 1, the proposed PIC algorithm has superior compression ratio performance than the standard gzip text compressor for all proteins over 20,000 atoms in size. This is seen visually in figure 5, as all except two points belonging to the two proteins with the fewest number of atoms lie above the diagonal, the region where PIC has better compression ratio performance. In figure 5, the gzip compression ratio decays while PIC’s compression ratio increases with atom count. Furthermore, unlike most compression algorithms, we can visually inspect the transformed image because it is itself a projection mapping of the original 3D structure. In figures 2 and 3, we show the PIC outputted images. For easier viewing, these images are inverted, five-fold contrast enhanced versions of the actual images outputted by the PIC compressor.

These results were obtained by running the *PIC*.*py* script in the command terminal with the “-e” option. These experiments were ran on a Ubuntu 20.04.4 LTS machine with an AMD Ryzen Threadripper 3970X 32-Core Processor and 256 GB of memory in single-thread mode without parallelization. However, the code has also been tested on an Apple MacBook Pro with a 3.5 GHz dual-core processor and 16 GB of memory, with comparable results. Thus, the code can run nearly as well on personal laptops.

### 3.2 Full PDB/mmCIF compression benchmark

Although PIC is designed as a prototype to showcase image-centric compression and thus only compressed the atomic point clouds, it is still instructive to compare against full compression software, such as MMTF, BCIF, and Foldcomp. In order to create a fair comparison, the total metadata space also needs to be included when comparing PIC—as such, we decided to use use MMTF to compress only the metadata, and then add that size to the size of the PIC image output. This is of course impractical for use as a compressor, but simply serves to level the playing field.

In Table 2, we compare the same benchmark proteins as in Table 1 with original PDB size, BCIF, MMTF, MMTF-lossy, PIC+MMTF-meta, and Foldcomp. MMTF-lossy notably both decreases precision to tenth of an Angstrom (same as PIC), but also only stores the C-alpha atoms, which allows them to take the least space at the cost of not storing all atoms. We were only able to get Foldcomp to work on one of our proteins, 4v60, because most of our benchmark proteins had discontinuous chains, which is not supported by Foldcomp, and several of the other proteins caused segfaults. However, on 4v60, Foldcomp does substantially better than PIC or any of the other compressors other than MMTF-lossy.

**Table 2:**
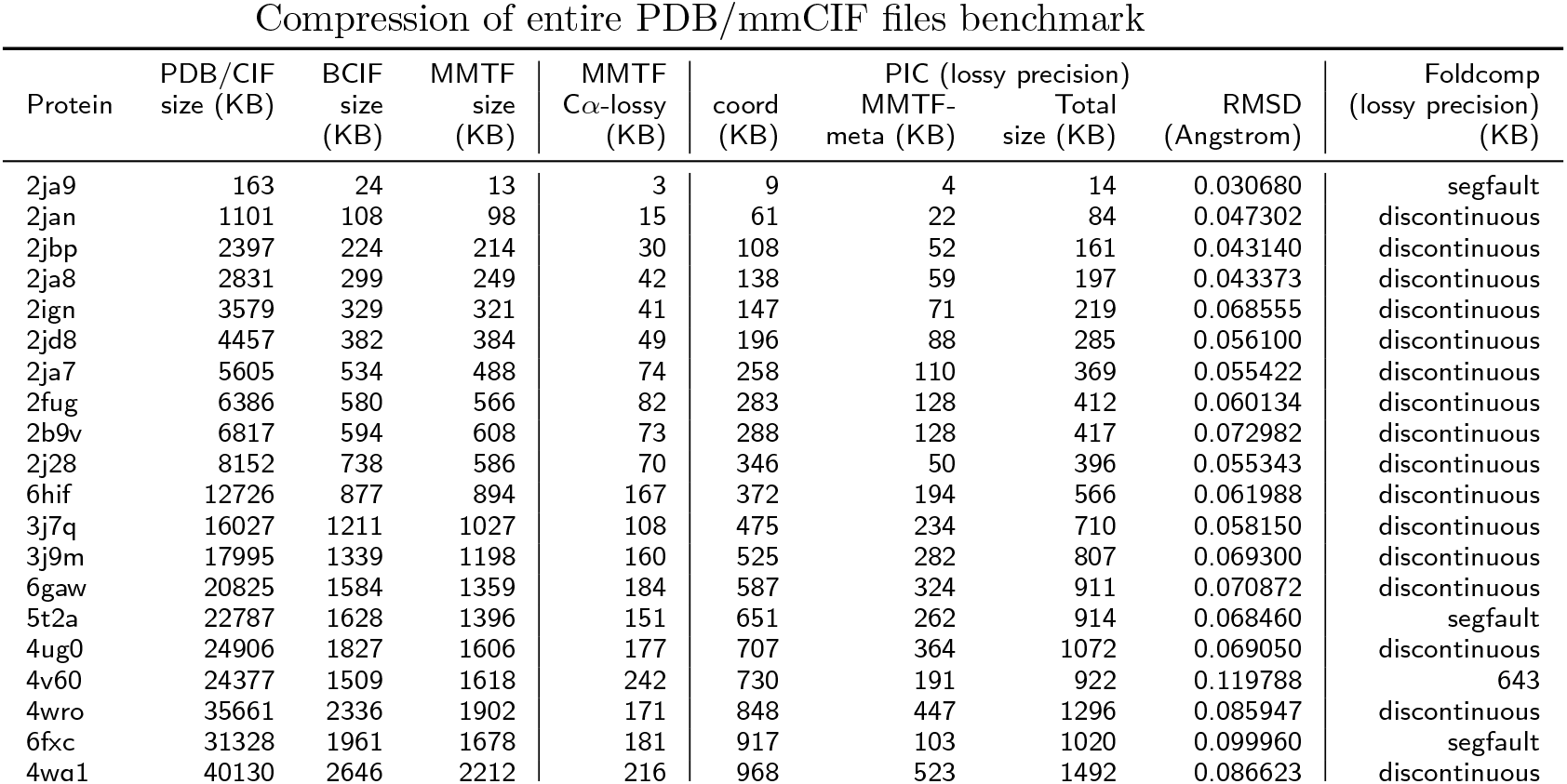
Actual compressed file sizes. We compare PIC, MMTF, Foldcomp, and BCIF formats. However, because PIC does not compress atomic metadata, we compressed the metadata-only with MMTF and then added that to the PNG sizes from PIC. All uses of MMTF were followed-up with standard Gzip compression on the MMTF file, as is standard, whereas BCIF is already a fully compressed file format. All sizes are in kilobytes (1000 *×* bytes). We also include the RMSD for PIC coordinates here. Lastly, unfortunately, most of the proteins we chose for our benchmark were discontinuous, or had other quirks, so Foldcomp was unable to handle them, but Foldcomp does substantially better oin 4v60 than all but MMTF-reduced, which not only decreases precision but also only keeps the alpha carbons.

## 4 Discussion

As expected, as atom count increases, more images are populated by PIC and more of the image space of the constructed images is used. In addition to the increased data load in only a slightly larger image width wise, this is due to an increased number of collisions as atom count increases. This causes the use of pointers, increasing the average number of bits used per data point, and, when no alternate location can be found in the current image, the population of a new image, increasing the image count.

Furthermore, as noted in the respective figures, our prototype PIC implementation is not optimized for speed. It is not intended as a drop-in replacement for gzip or MMTF, but is instead meant to show that image-centric compression of protein atomic point clouds can provide significant space savings. The Python implementation takes on the order of a few minutes for a single compression/decompression, which is significantly slower than the order of seconds for gzip compression.

Other values of *ε* investigated include {1.25, 5, 10}. Only the results for *ε* = 2.5 are shown as this value produced the best compression ratios. As stated in section 2.1, higher *ε* increased image sizes and consequently decreased the compression ratios. Setting *ε* = 1.25 increased compression times as collisions increased due to the decreased image size and alternate mapping locations needed to be considered. Compression ratios also decreased slightly as gains in the compression ratios from decreased image sizes were overcame by the additional use of pointers and higher number of images generated by the PIC algorithm. Importantly, in this prototype study, we have given results from only a single set of parameters for all sizes of proteins for principled benchmarking. In future work, it may be preferable to set parameters dynamically for each protein and to store them, as in standard practice in many file compression formats. This will play a greater role as the structures of more complex proteins are deconstructed, stored in databases, and transmitted amongst researchers. Further, as the PIC algorithm leverages the standard and widely used PNG image compressor, the algorithm can be easily implemented on a variety of platforms and systems.

Although we ran benchmarks, the comparisons against MMTF, BCIF, and Foldcomp were not especially informative for a couple of reasons. First, PIC does not compress metadata, so we had to use MMTF for that portion to create reasonable comparisons. Second, different tools made different design choices about what to focus on: MMTF-lossy also only stores the C-alpha atoms, rather than all atoms as PIC and Foldcomp do, and Foldcomp is not designed for discontinuous chains, which were common in our benchmark data set. Still, it does seem that if only C-alpha atomic coordinates are needed, MMTF-lossy is better, and where Foldcomp works, it is also better.

## 5 Conclusion

In this paper, we have introduced PIC, an new compression algorithm that leverages positional encoding techniques and the well-developed, widely available PNG image compressor to encode and compress structural protein data in PDB and mmCIF files. The algorithm encodes two of the three dimensions of an atomic coordinate from the point cloud stored in the file to a position in the image space and stores the remaining dimension in pixels’ intensity values around that location. The resulting image is then compressed with the lossless image compressor PNG. We showed PIC has a compression ratio superior of that of gzip for proteins with more than 20,000 atoms, and improves with the size of the protein being compressed, reaching up to 37.4% on the proteins we examined. Although as of September 2023, only 6.7% of the structures in the RCSB PDB are over 20,000 atoms, they represent at least 21% of the database in terms of total atom count, so PIC has fair applicability. The improvement in compression is also orthogonal to the lossy storage of atomic coordinates to a precision of only a tenth of an angstrom, as we compared against gzip on files with that precision level.

More important than just providing a prototype, we demonstrate in this paper that the paradigm of image-centric compression is superior in efficacy than simply applying a standard sequential compressor to the atomic point clouds. This result is consistent with examples from point-cloud compression in LIDAR imaging, read-reordering for NGS sequence compression, and also the recent Foldcomp compressor which stores internal angles instead. Importantly, this improvement in compression ratios persists even though we store the necessary metadata to undo any atom-reorderings; thus, the only lossy portion of PIC is in coordinate positions. Still, we would recommend that the ordering information be entirely discarded, as it is for LIDAR and read re-ordering—we only kept all of that information to ensure that we performed a fair comparison in our benchmarks. Were we to discard that information, PIC’s benchmark results would be even stronger.

We do note that as a prototype implementation, our runtimes and file formats are not suitable for everyday use, but our hope is that future compression algorithms will be designed with our findings in mind. Ultimately, we hope that this study points the way for future image-centric (or more generally structure-aware) compression of protein structures. Indeed, the contemporaneous Foldcomp [22] makes use of internal bond angles and torsions in protein compression, which is a different means of exploiting the 3D structure than our image-centric approach, and that shows greater promise even than PIC.

The PIC algorithm itself, if reimplemented in a faster language, is certainly competitive on compression ratios already, and furthermore is easy to implement be-cause the PNG image format is already implemented on many platforms. However, we mostly envision that image-centric compression will simply form a part of other more complex compression methods. As structural protein files with increasing complexity are deconstructed, added to databases, and transmitted amongst researchers, targeted compression techniques will become ever more necessary.

## Acknowledgements

We would like to thank Jim Shaw, Ziye Tao, and Alex Leighton for their thought-provoking questions that inspired parts of this work.

## Funding

This work was supported by the Ontario Graduate Scholarship Program. We acknowledge the support of the Natural Sciences and Engineering Research Council of Canada (NSERC), (NSERC grant RGPIN-2022-03074), and the DND/NSERC Discovery Grant Supplement for 2022.

### Abbreviations

PDB: Protein Data Bank
mmCIF: macromolecular Crystallographic Information File
LIDAR: light detection and ranging

## Availability of data and materials

Project name: PIC Compression and Decompression Prototype

Project home page: https://github.com/lukestaniscia/PIC

Operating system(s): Platform independent

Programming language: Python

Other requirements: Python 3.0 or higher License: MIT License

Any restrictions to use by non-academics: No

## Ethics approval and consent to participate

Not applicable.

## Consent for publication

Not applicable.

## Authors’ contributions

L.S. and Y.W.Y. jointly conceived of the study. L.S. implemented most of the code, under Y.W.Y.’s oversight. Both authors were fully involved in writing the manuscript.

## References

1. Ramachandran, G.: Protein structure and crystallography. Science, 288–291 (1963)

2. Ilari, A., Savino, C.: Protein structure determination by x-ray crystallography. Bioinformatics, 63–87 (2008)

3. Rose, P.W., Prlić, A., Altunkaya, A., Bi, C., Bradley, A.R., Christie, C.H., Costanzo, L.D., Duarte, J.M., Dutta, S., Feng, Z., et al.: The rcsb protein data bank: integrative view of protein, gene and 3d structural information. Nucleic acids research, 1000 (2016)

4. Berman, H.M., Kleywegt, G.J., Nakamura, H., Markley, J.L.: The protein data bank at 40: reflecting on the past to prepare for the future. Structure 20(3), 391–396 (2012)

5. Pearson, W.R.: Using the fasta program to search protein and dna sequence databases. In: Computer Analysis of Sequence Data, pp. 307–331. Springer, ??? (1994)

6. Westbrook, J.D., Fitzgerald, P.M.: The pdb format, mmcif formats, and other data formats. Structural bioinformatics 44, 159–179 (2003)

7. Varadi, M., Anyango, S., Deshpande, M., Nair, S., Natassia, C., Yordanova, G., Yuan, D., Stroe, O., Wood, G., Laydon, A., et al.: Alphafold protein structure database: massively expanding the structural coverage of protein-sequence space with high-accuracy models. Nucleic acids research 50(D1), 439–444 (2022)

8. Fritz, M.H.-Y., Leinonen, R., Cochrane, G., Birney, E.: Efficient storage of high throughput dna sequencing data using reference-based compression. Genome research 21(5), 734–740 (2011)

9. Daniels, N.M., Gallant, A., Peng, J., Cowen, L.J., Baym, M., Berger, B.: Compressive genomics for protein databases. Bioinformatics 29(13), 283–290 (2013)

10. Yu, Y.W., Yorukoglu, D., Peng, J., Berger, B.: Quality score compression improves genotyping accuracy. Nature biotechnology 33(3), 240–243 (2015)

11. Hernaez, M., Pavlichin, D., Weissman, T., Ochoa, I.: Genomic data compression. Annual Review of Biomedical Data Science 2, 19–37 (2019)

12. Hategan, A., Tabus, I.: Protein is compressible. In: Proceedings of the 6th Nordic Signal Processing Symposium, 2004. NORSIG 2004., pp. 192–195 (2004). IEEE

13. Pinho, A.J., Pratas, D.: MFCompress: a compression tool for FASTA and multi-FASTA data. Bioinformatics 30(1), 117–118 (2013). doi:10.1093/bioinformatics/btt594. https://academic.oup.com/bioinformatics/article-pdf/30/1/117/17124293/btt594.pdf

14. Deorowicz, S., Walczyszyn, J., Debudaj-Grabysz, A.: CoMSA: compression of protein multiple sequence alignment files. Bioinformatics 35(2), 227–234 (2018). doi:10.1093/bioinformatics/bty619. https://academic.oup.com/bioinformatics/article-pdf/35/2/227/27497129/bty619supplementarymaterial.pdf

15. Kryukov, K., Ueda, M.T., Nakagawa, S., Imanishi, T.: Nucleotide Archival Format (NAF) enables efficient lossless reference-free compression of DNA sequences. Bioinformatics 35(19), 3826–3828 (2019). doi:10.1093/bioinformatics/btz144. https://academic.oup.com/bioinformatics/article-pdf/35/19/3826/30061651/btz144.pdf

16. Cox, A.J., Bauer, M.J., Jakobi, T., Rosone, G.: Large-scale compression of genomic sequence databases with the burrows–wheeler transform. Bioinformatics 28(11), 1415–1419 (2012)

17. Hach, F., Numanagić, I., Alkan, C., Sahinalp, S.C.: Scalce: boosting sequence compression algorithms using locally consistent encoding. Bioinformatics 28(23), 3051–3057 (2012)

18. Patro, R., Kingsford, C.: Data-dependent bucketing improves reference-free compression of sequencing reads. Bioinformatics 31(17), 2770–2777 (2015)

19. Goodsell, D.S.: PDB101: Learn: Guide to Understanding PDB Data: Introduction to PDB Data (n.d.). https://pdb101.rcsb.org/learn/guide-to-understanding-pdb-data/introduction

20. Bradley, A.R., Rose, A.S., Pavelka, A., Valasatava, Y., Duarte, J.M., Prlić, A., Rose, P.W.: Mmtf—an efficient file format for the transmission, visualization, and analysis of macromolecular structures. PLoS computational biology 13(6), 1005575 (2017)

21. Sehnal, D., Bittrich, S., Velankar, S., Koča, J., Svobodová, R., Burley, S.K., Rose, A.S.: Binarycif and ciftools—lightweight, efficient and extensible macromolecular data management. PLoS computational biology 16(10), 1008247 (2020)

22. Kim, H., Mirdita, M., Steinegger, M.: Foldcomp: a library and format for compressing and indexing large protein structure sets. bioRxiv (2022)

23. Valasatava, Y., Bradley, A.R., Rose, A.S., Duarte, J.M., Prlić, A., Rose, P.W.: Towards an efficient compression of 3d coordinates of macromolecular structures. PLOS ONE 12(3), 1–15 (2017). doi:10.1371/journal.pone.0174846

24. Deutsch, P., et al.: Gzip file format specification version 4.3. RFC Editor (1996)

25. Alakuijala, J., Farruggia, A., Ferragina, P., Kliuchnikov, E., Obryk, R., Szabadka, Z., Vandevenne, L.: Brotli: A general-purpose data compressor. ACM Transactions on Information Systems (TOIS) 37(1), 1–30 (2018)

26. Houshiar, H., Nüchter, A.: 3d point cloud compression using conventional image compression for efficient data transmission. In: 2015 XXV International Conference on Information, Communication and Automation Technologies (ICAT), pp. 1–8 (2015). doi:10.1109/ICAT.2015.7340499

27. Divne, C.: 2IGN: Crystal structure of recombinant pyranose 2-oxidase H167A mutant (2006). https://www.rcsb.org/structure/2IGN

28. Kato, K., Zhou, Y., Tanaka, H., Yao, M., Yamashita, M., Tsukihara, T.: 4V60: The structure of rat liver vault at 3.5 angstrom resolution (2014). https://www.rcsb.org/structure/4V60

